# Predicting Differential Protein-Protein Interactions

**DOI:** 10.64898/2026.10.29.762715

**Authors:** Aram Papazian, Volkhard Helms

## Abstract

**Motivation:** Analysis of context-specific protein-protein interactions (PPIs) provides deeper insight into protein function, revealing how cellular pathway components interact in certain perturbation conditions. However, identifying such interactions experimentally often requires combining affinity purification with mass spectrometry, proving costly and time-consuming. On the other hand, it has been shown that an increase in protein abundance does not always correspond to an increase in protein interaction, making it an unsuitable surrogate for interaction studies.

**Results:** To address this, a random forest model was created using PPI-specific MINT embeddings and condition-specific mass spectrometry protein abundance information to predict differential interactions across two experimental conditions. The model was trained and tested on validated context-specific PPIs from eight distinct cell-lines, demonstrating proficiency in predicting differential PPIs for both novel proteins and cell lines. To our knowledge, we provide the first computational model that predicts differential PPIs between experimental conditions.

**Availability and implementation:** The code is available on GitHub and backed up on Zenodo (DOI: 10.5281/zenodo.23758127). The model is freely available on Zenodo (DOI: 10.5281/zenodo.23757927).

## Introduction

Proteins are essential components in nearly all biological cellular processes, including cellular transport and signaling. While some carry out biological roles independently, most proteins function while bound to other proteins. Thus, protein functionality is often dependent on the efficacy of such interactions [1]. Some assemblies remain stably bound over prolonged periods of time, such as ribosomes or RNA polymerases. Other protein complexes, such as those implied in signaling cascades or redox reactions, bind reversibly and sustain short-lived interactions. High-throughput techniques, such as tandem affinity purification and mass spectrometry (TAP-MS), have been established as standard techniques for generating proteome-wide interactomes in both *Homo sapiens* and *Saccharomyces* models. Data resulting from these techniques are compiled into specialized repositories, such as BioGRID or Mentha [2–3]. Remarkably, few contemporary studies report cell-specific and condition-specific protein-protein interactions (PPIs), which would depict a context-specific interactome [4–6].

When a PPI is present in one condition but absent in another, it is considered a differential interaction, often attributed to post-translational modifications [7]. Differential interactions are commonly detected by techniques combining affinity purification with mass spectrometry (AP-MS), both of which entail lengthy, resource-heavy protocols. For example, Huttlin et al. [4] incorporated results from 15,650 pull-down experiments and utilized AP-MS, CompPASS [8–9], and CompPASS-Plus [10] to identify PPIs in HEK293T and HCT116 cell lines. Moreover, Kim et al. [5] combined AP-MS, CompPASS, and SAINTexpress [11] to identify PPIs across the three breast cell lines MCF7, MDA231, and MCF10A. Using the same approach, Swaney et al. [6] were able to identify PPIs in the head and neck cancer cell lines SCC25, CAL33, and HET1A. Unlike [4], [5] and [6] focused on bait proteins that are known to be altered under their respective conditions. Hence, smaller numbers of identified interactions were reported.

With the use of standard mass spectrometry (MS), Kim et al. [5] reported that differential protein abundances can result in cell type-specific enriched PPIs, but they also detected many cases with the opposite effect where a gain in interaction was observed despite a concomitant decrease in protein abundance. Therefore, differential abundances alone cannot be used to reliably determine PPIs. Verifying co-expression is another approach to resolve PPIs, proving especially useful for permanent interactions in which complex members are known to co-express [12–13]. With the advent of pretrained protein language models (PLM), such as evolutionary scale models [14–15], several sequence-based deep learning (DL) techniques have been recently developed to predict PPIs. Two prominent PLM models based on the data-leakage free Bernett et al. dataset [16] are TUnA [17] and PLM-interact [18]. Both models utilized pretrained ESM2 [14] embeddings and diverse inter-protein attention mechanisms to achieve the best performance on predicting PPIs. However, one limitation of these models is that their predictions do not differ between contexts, but they can act as signal for interaction strength.

In our study, a random forest (RF) model was trained on experimentally validated context-specific PPIs across eight distinct human cancer cell lines [4–6]. The model inputs cell line-specific MS intensities and sequence-based protein interaction embeddings to predict differential interactions between two experimental conditions. The model showed formidable performance on both a sequence similarity-based and multiple external cell line test sets. We believe that our model will be useful in identifying key dysregulated PPIs that lead to disease development and to our knowledge this is the first computational tool that is able to predict differential interactions between two conditions.

## Methods and Materials

### AP-MS Data

Cell line-specific, experimentally validated sets of PPIs and non-PPIs were retrieved from three studies, spanning eight human cancer cell lines. In particular, HEK293T and HCT116 cell line PPIs were derived from Supplementary Table 1 of Huttlin et. al [4]; an interaction was considered present if it was within the top 5% of the cell line’s candidate interactions, as annotated in Huttlin et. al [4] and absent otherwise. MCF7, MDA231, and MCF10A cell line interactions were retrieved from Supplementary Table 3 of Kim et. al [5]; in this case, an interaction was considered present if it possessed a PPI score greater than 0.9 and considered absent otherwise. This threshold was established by the authors to define high-confidence protein interactions [5]. CAL33, SCC25, and HET1A cell line PPIs were acquired from Supplementary Table 2 of Swaney et. al [6], with interactions considered present if annotated as a high-confident PPI [6] and absent otherwise. In all three studies, only wild-type PPIs were considered.

### Mass Spectrometry Data

As context-specific data, MS abundance values were taken into account as detailed within these three studies [4–6]. Huttlin et. al [4] reported five replicates per protein, while Kim et. al and Swaney et. al [5–6] used only three. Further cell line-specific protein abundance data were retrieved from Gonçalves et. al [19], which provided six replicates per protein for lines HCT116, CAL33, SCC25, MCF7 and MDA231. Incorporating these supplementary abundance data allowed for examination of detected interactions with respect to external MS intensity values. For all cell lines, preprocessing was carried out by removing proteins in the MS datasets possessing less than 70% valid values. Then, imputation of the remaining missing values was performed using KNN, with k=10. All replicate columns underwent median normalization. For the HEK293T, HET1A and MCF10A lines, these steps were carried out only on original MS data provided by each study [4–6]. For the remaining cell lines, preprocessing was performed separately for the original and external MS data [19].

Ultimately, cell line-specific interactions were filtered to include only MS-detectable proteins that met the filtering criteria. The total remaining present and absent PPIs for each cell line per source, original or external, are listed in Table 1.

**Table 1.** Quantity of remaining experimentally verified present and absent PPIs, based on detected MS intensities per source. “_Ori” indicates MS intensities retrieved from the original studies, while “_Ext” signifies external intensities supplemented from Gonçalves et. al [19].

| Cell Line | Present Interactions | Absent Interactions |
| --- | --- | --- |
| HEK293T_Ori | 55,350 | 13,649 |
| HCT116_Ori | 35,000 | 33,999 |
| HCT116_Ext | 1533 | 1034 |
| MCF7_Ori | 100 | 26,938 |
| MCF7_Ext | 74 | 8,326 |
| MDA213_Ori | 115 | 20,075 |
| MDA213_Ext | 110 | 13,830 |
| MCF10A_Ori | 58 | 9,896 |
| CAL33_Ori | 127 | 34,031 |
| CAL33_Ext | 25 | 10,785 |
| SCC25_Ori | 80 | 21,462 |
| SCC25_Ext | 9 | 4,786 |
| HET1A_Ori | 166 | 35,566 |

### Differential PPIs

To annotate differential PPIs between cell lines, each pair of cell lines was analyzed separately. If an interaction is shared between two cell lines and exhibits the same present/absent label, it is labelled as “conserved”. However, when the interaction is shared but has different labels, it is considered “differential”. Cell line comparisons were performed only if both cell lines possessed MS intensities that were identically processed, due to similar protein intensity quantification techniques used. For example, the external MS data from the HCT116, MDA231, MCF7, CAL33, and SCC25 obtained from Gonçalves et. al were compared against each other [19]. Though, for the HEK293T line, which only had MS data featured in Huttlin et. al [4], the cell line could only be compared to the original HCT116 MS intensities [4]. Similarly, the MDA231, MCF7, MCF10A, CAL33, SCC25, and HET1A cell lines were compared against each other with respect to MS intensities [5–6]. The quantities of conserved and differential interactions are listed in Table 2. The leftmost column displays the pairs of cell lines compared.

**Table 2.** Total conserved and differential interactions between cell line pairs. “_Ori” designates MS intensities retrieved from the original studies, while “_Ext” indicates those retrieved from Gonçalves et. al [19].

| Cell Line Pair | Conserved Interactions | Differential Interactions |
| --- | --- | --- |
| CAL_Ext / HCT_Ext | - | 1 |
| CAL_Ext / MDA_Ext | 2,281 | 6 |
| CAL_Ext / SCC_Ext | 2,166 | - |
| CAL_Ori / HET_Ori | 25,330 | - |
| CAL_Ori / MCF10A_Ori | 1,345 | 15 |
| CAL_Ori / MCF7_Ori | 1251 | 10 |
| CAL_Ori / MDA_Ori | 4,322 | 20 |
| CAL_Ori / SCC_Ori | 18,931 | - |
| HCT_Ori / HEK_Ori | 16,874 | 36,254 |
| HCT_Ext / MCF7_Ext | 2 | 3 |
| HCT_Ext / MDA_Ext | 6 | 6 |
| HCT_Ext / SCC_Ext | 1 | 1 |
| HET_Ori / MCF10A_Ori | 1,953 | 57 |
| HET_Ori / MCF7_Ori | 1,966 | 33 |
| HET_Ori / MDA_Ori | 4,140 | 18 |
| HET_Ori / SCC_Ori | 17,659 | - |
| MCF10A_Ori / MCF7_Ori | 4,721 | 40 |
| MCF10A_Ori / MDA_Ori | 5,131 | 62 |
| MCF10A_Ori / SCC_Ori | 1,253 | 16 |
| MCF7_Ext / MDA_Ext | 3,972 | 44 |
| MCF7_Ext / SCC_Ext | 3 | - |
| MCF7_Ori / MDA_Ori | 10,648 | 59 |
| MCF7_Ori / SCC_Ori | 1,100 | 12 |
| MDA_Ext / SCC_Ext | 894 | 1 |
| MDA_Ori / SCC_Ori | 2,542 | 15 |

### MS-Based Features

Furthermore, MS protein intensities were implemented as context-specific features. For each interaction, its pair of protein intensity rows were aligned across both cell lines, as visualized in Figure 1. Resulting from this alignment, a set of features were derived, as detailed in Table 3.

**Figure 1.**
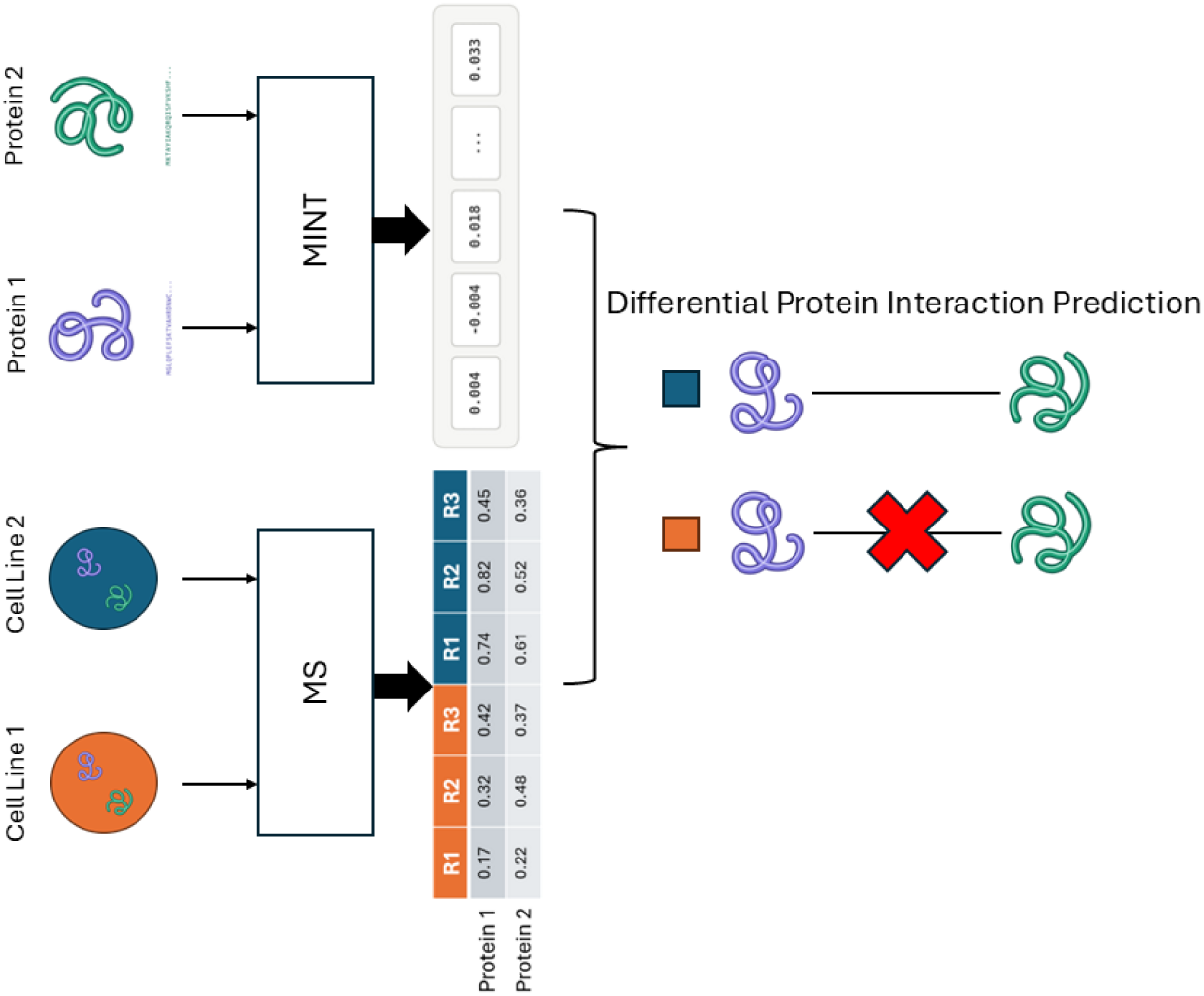
Combining MS abundance features with MINT PPI interaction embeddings to predict differential PPIs between two conditions.

**Table 3.**
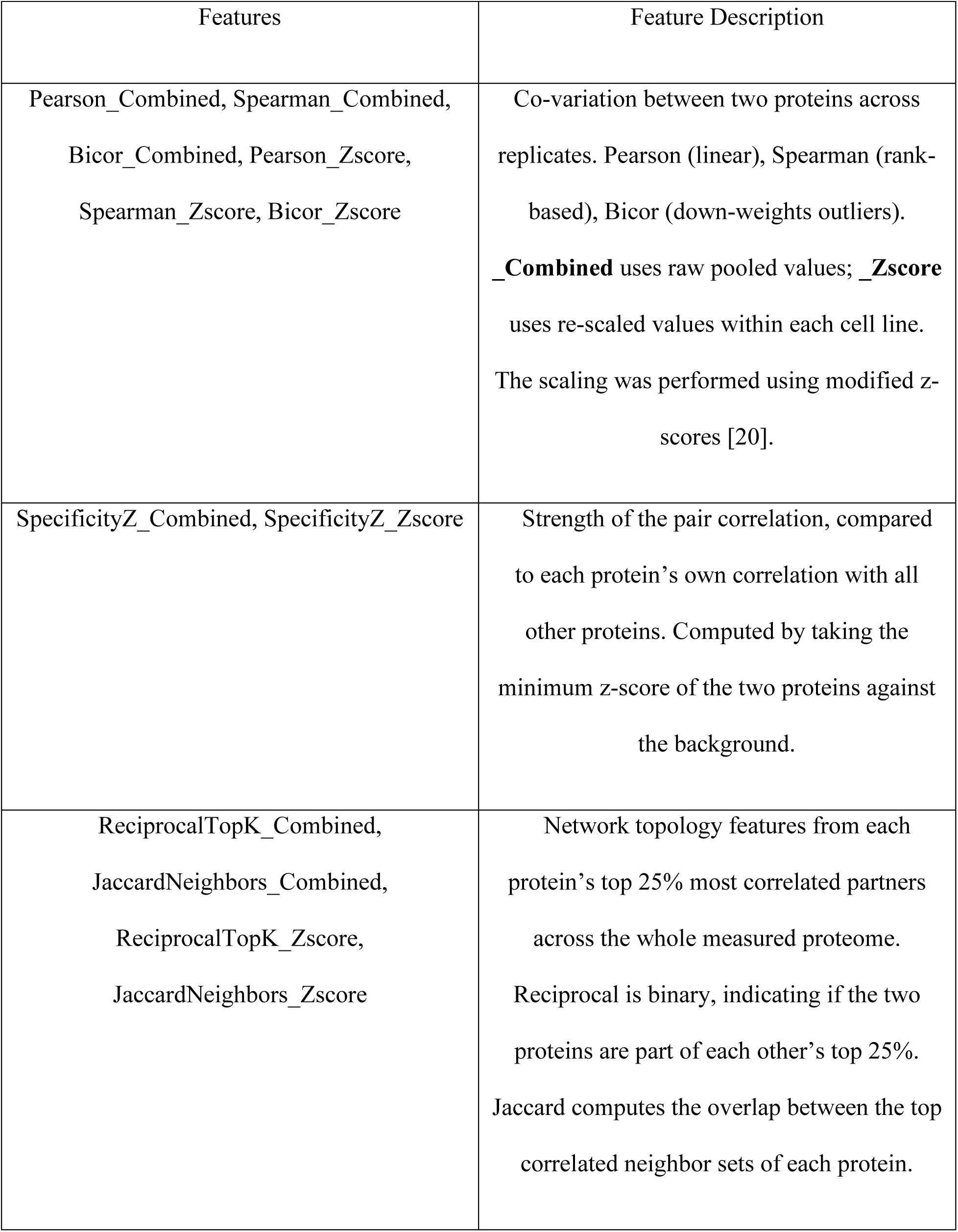
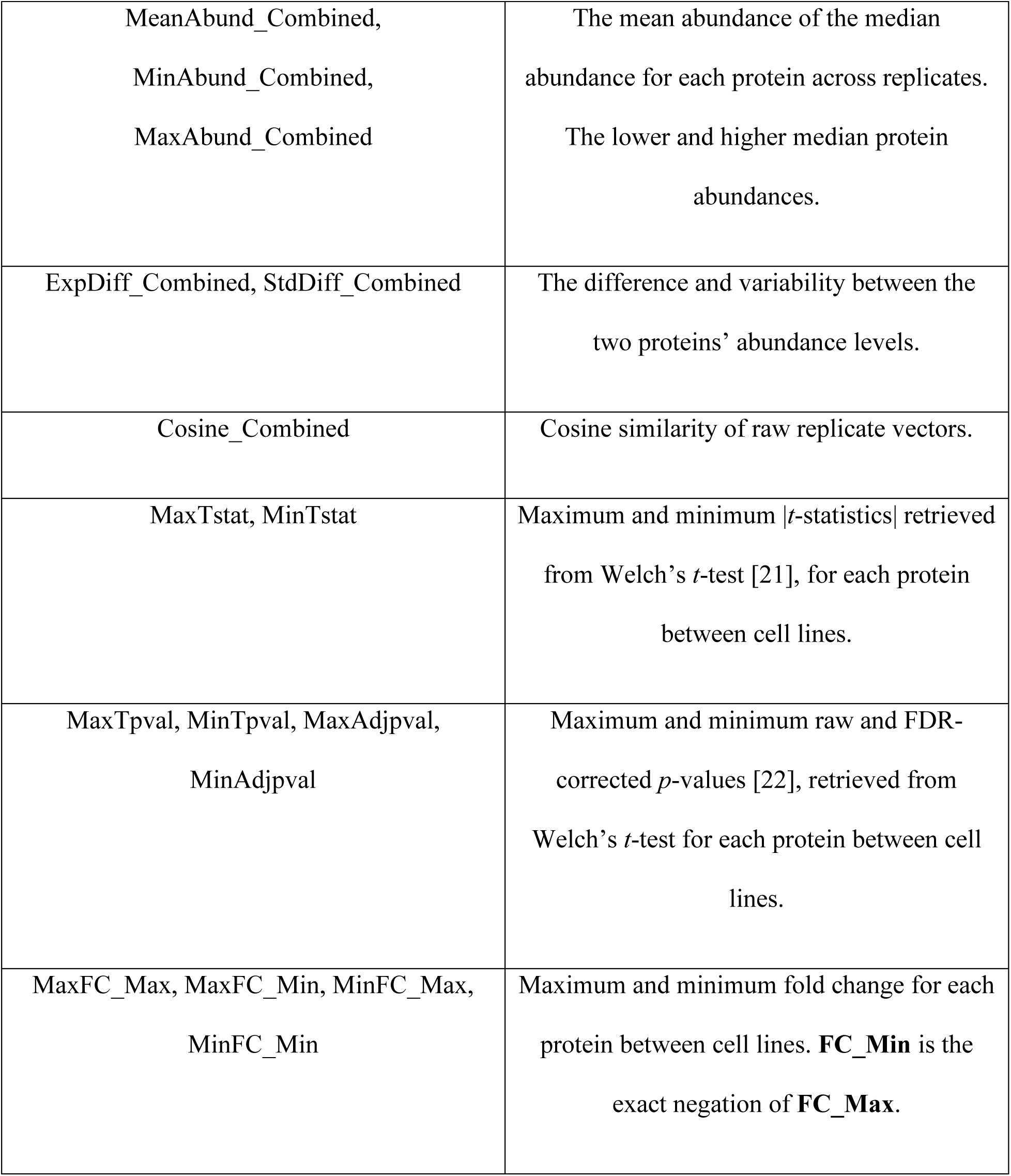
Features derived from MS protein expression data between pairs of cell lines. Maximum and minimum values were taken to avoid order-specific features. See Supplement for more details.

### Sequence-Based Features

Protein sequence information can be used to determine whether two proteins interact. Several well-established sequence-based models have been developed to accurately predict PPIs, though not taking experimental conditions into account [14, 17, 18]. For this study, embeddings from MINT [23], a PLM designed to model sets of interacting proteins in a contextual scalable manner, were used; see Figure 1. MINT was derived by conducting unsupervised training on a large, curated PPI dataset compiled from the STRING database and was shown to outperform existing PLMs in diverse PPI tasks [24].

MINT outputs a sequence embedding for each protein separately. To obtain a PPI-level embedding for a pair of proteins, both protein sequence embeddings were mean pooled, resulting in a total of 1280 sequence-based features.

### Random Forest

Based on the aforementioned features, using *sklearn version 1.7.2* [25], a random forest classifier was trained to predict differential interactions between two experimental conditions. Table 2 clearly shows that there exists both an interaction type label and cell line pair imbalance. To mitigate this, a sample weighting scheme was implemented:

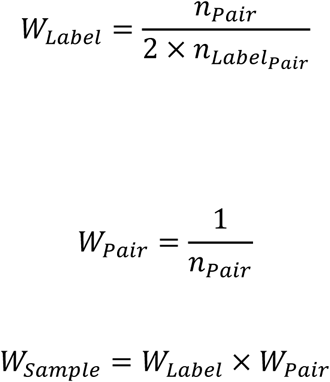

Here, *n_Pair_* represents the total number of interactions within a certain cell line pair. If there exist fewer than 20 interactions within a pair, *n_Pair_* is set to 20 by default. *n_LabelPair_* represents the number of conserved or differential interactions within a pair, while *W_Label_*, *W_Pair_* and *W_Sample_* represent the computed label, pair, and sample weights, respectively. The samples weights are then normalized by the mean sample weight.

The sample weights affect the random forest by adjusting the way in which Gini impurity [26] is computed. Here, label frequencies were accumulated at each node as weighted sums, rather than raw counts. Thus, split selection is scored against weighted label proportions, rather than observed label proportions. Additionally, the same weighted proportions determine each leaf’s predicted probability, which is the weighted fraction of positives, signifying differential interactions, as opposed to the raw fraction.

### Data Splitting

To benchmark the performance of the weighted random forest, two data splitting criteria were formulated. First, a sequence similarity-based split was performed using DataSail [27] and MMseqs2 [28], with a sequence similarity threshold of 0.3 and coverage of 0.8. Interactions were kept only in cases in which both proteins were assigned the same splitting annotation, resulting in a single optimal split. With respect to cell lines, the second splitting criterion tested the model on interactions belonging to a pair of cell lines not present in the training set. For example, in the case “MCF7_Ori / MDA_Ori” is tested, all cell line pairs involving either MCF7 or MDA are removed from the training set. The test sets aim to mitigate data leakage and determine if the model generalizes predictions for both unseen proteins and cell lines.

## Results

### Model Performance

To test whether our model can generalize to novel proteins and cell lines, the two aforementioned different splitting protocols were conducted. For the sequence similarity-based split, only interactions in which both proteins possess the same splitting annotation were kept. This resulted in a training set of 27,341 conserved and 13,565 differential interactions, as well as a test set consisting of 34,056 conserved and 5,820 differential interactions. The performances of regular and weighted random forest models are illustrated in Figure 2; here, the task of the model is to predict whether a certain PPI is differential (positive) or conserved (negative) between two cell lines.

**Figure 2.**
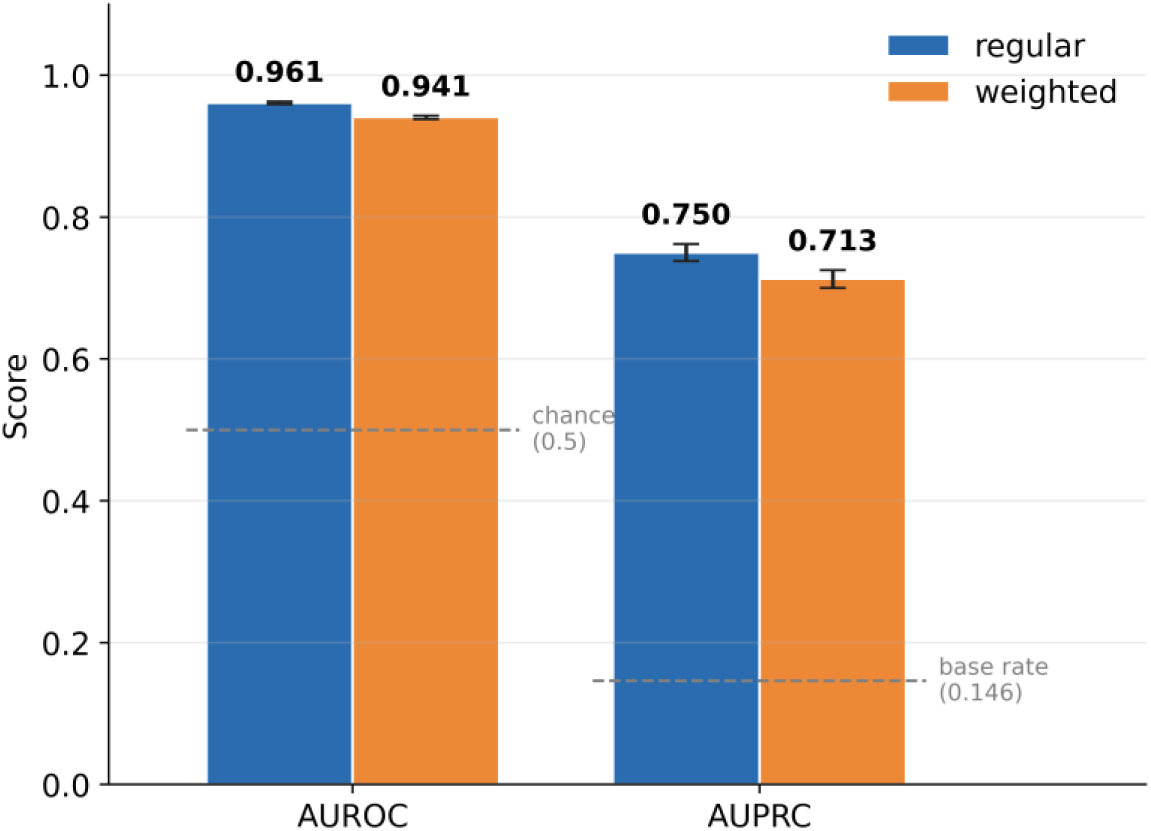
Performance of the regular and weighted RF models on the sequence similarity-based test set. The 95% confidence intervals were computed by stratified resampling of the test set to preserve the original label proportions.

It is evident that both RF models performed proficiently on this test set. Both models possessed ROC and PR AUC scores that substantially exceeded the baseline score. Weighting the samples slightly degraded model performance, as the weighted RF consistently achieved lower scores. This can be attributed to the weighting scheme balancing the influence of the different cell line pairs, as opposed to focusing on the pooled distribution of conserved and differential interactions across pairs.

To compare these models with respect to cell line pairs, using the aforementioned prediction task, the models were also tested across 14 distinct test sets; see Figure 3. Each test set was composed of a cell line pair not seen during training, verifying the models’ ability to generalize to unseen conditions.

**Figure 3.**
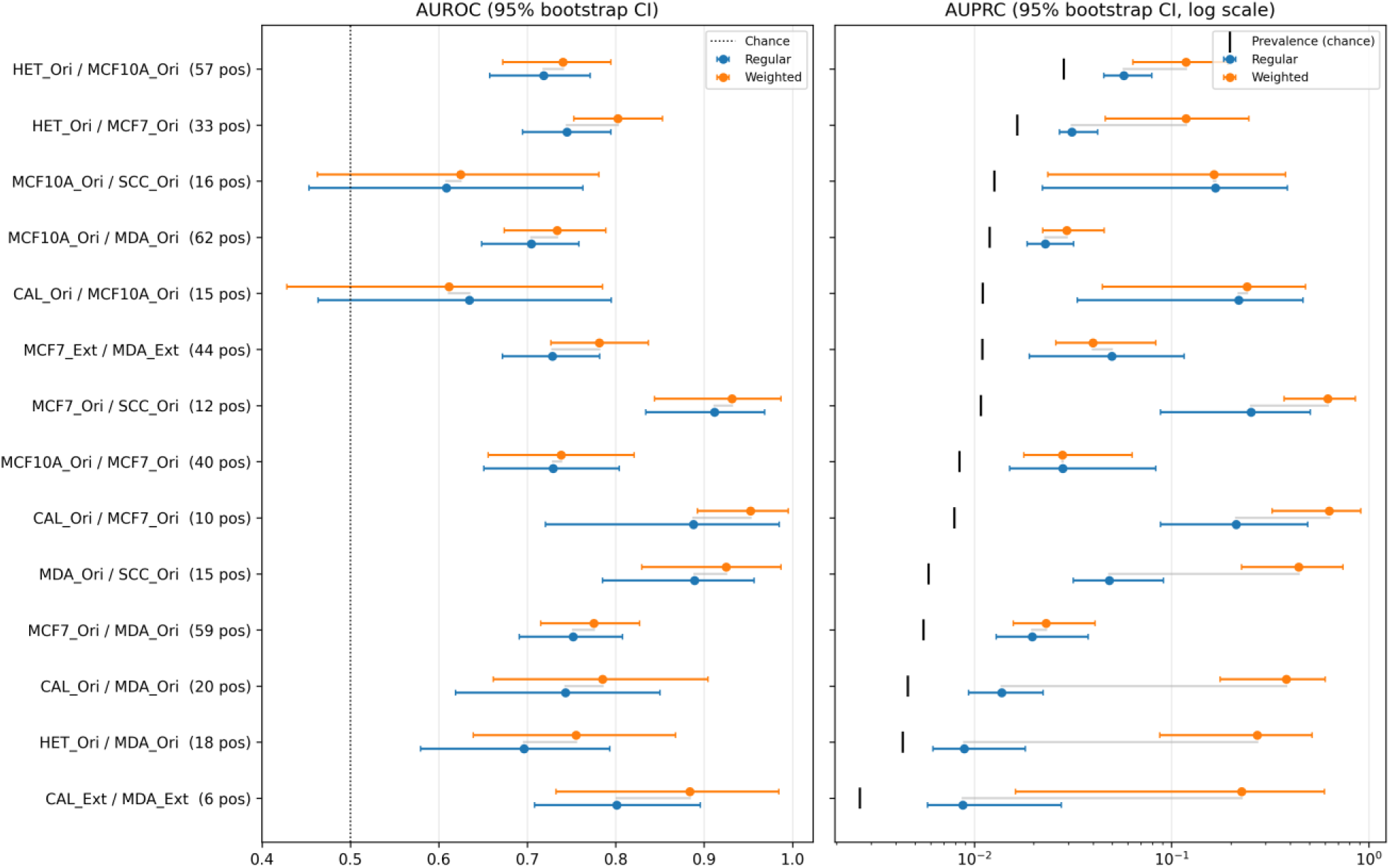
Performance of the weighted and regular RF models on 14 unseen cell line pair test sets. Cell line pairs are annotated with the number of differential interactions (pos) in each set. The prevalence of the differential interactions was used as a baseline measurement for AUPRC.

As observed in Figure 3, the weighted RF outperforms the regular RF in predicting differential interactions for external cell line pairs. Namely, in the AUROC panel to the left, it is observed that the weighted RF achieved higher scores in 13 out of the 14 test sets, with a mean AUROC score of 0.79±0.1, compared to a mean score of 0.75±0.09 for the regular RF. For AUPRC, the weighted RF attained higher scores in 11 out of the 14 test sets, with a mean AUPRC score of 0.24±0.21, compared to a mean score of 0.08±0.09 for the regular RF. The standard deviation scores of the AUPRC score range higher due to the varying prevalence of differential interactions within each test set.

### Differentiation of Conserved Interactions

To probe the model outputs further, the contribution of the MS and embedding features were analyzed separately, which was performed on the sequence similarity-based test set, filtered for PPIs available in STRING [24]. Out of the 39,876 possible PPIs, only 16,590 PPIs were found. For this analysis, the conserved interactions were separated into interactions that were either present or absent in both conditions. Hence, the remaining test set interactions consisted of 3,943 differential, 10,554 conserved absent (CA), and 2,093 conserved present (CP) interactions. The STRING combined scores for each interaction type were compared against the weighted RF, MS-only, and embeddings-only model predictions. Additionally, the PLM-interact [18] and TUnA [17] model outputs were visualized for the different interaction types; see Figure 4.

**Figure 4.**
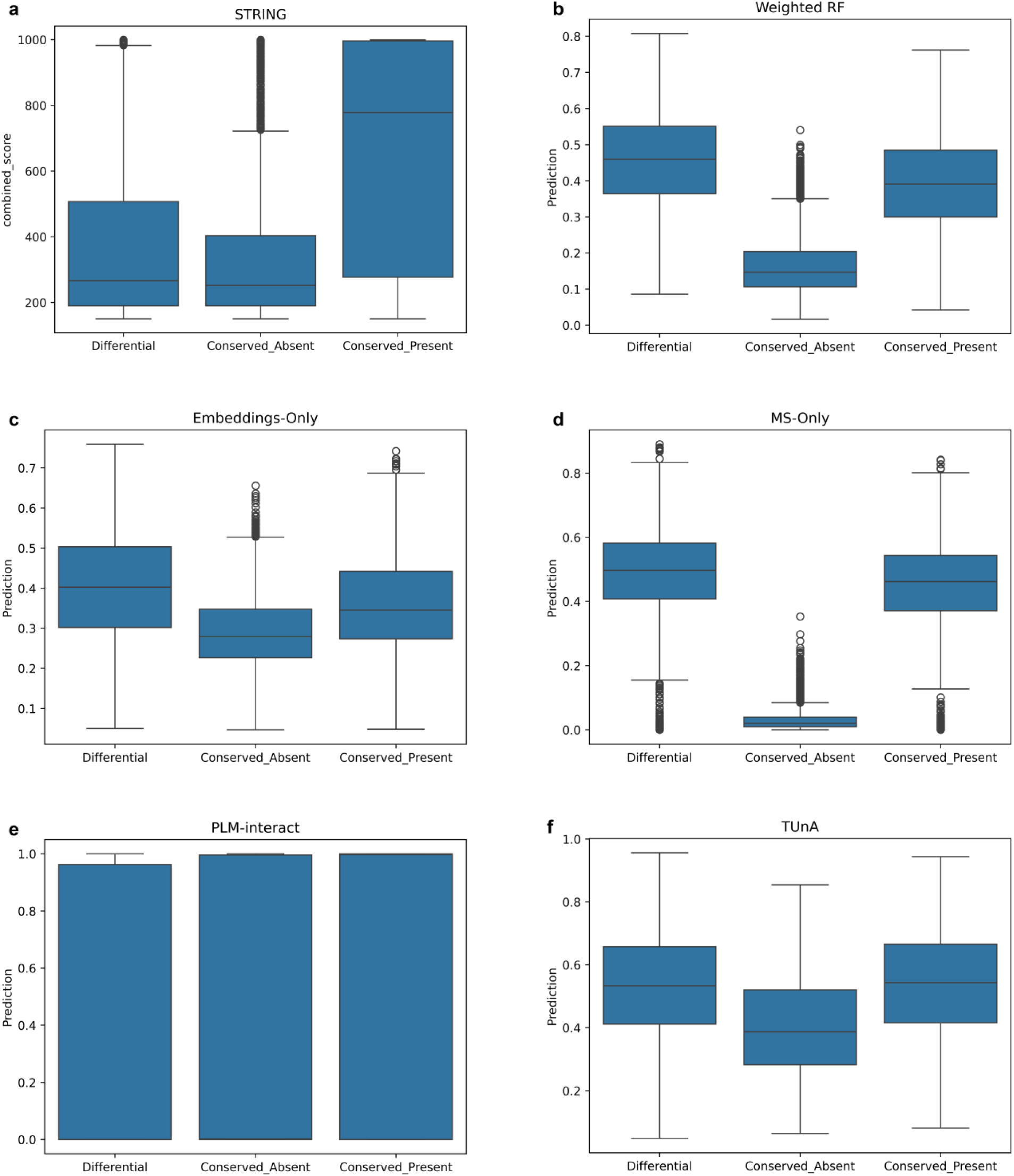
(**a**) STRING combined score distribution for the differential, conserved present, and conserved absent PPIs. (**b**) Weighted RF model predictions for the different interaction types. Weighted RF model predictions using either the MS-based features (**c**) or embeddings (**d**). PLM-interact (**e**) and TUnA (**f**) predictions for the different interaction types.

Figure 4a reveals that CP interactions tend to have the largest STRING combined scores, indicating that these PPIs are well annotated with high likelihoods of interaction. Though, STRING is unable to discern between differential and CA interactions, considering both types to have lower combined scores relative to CP interactions. The weighted RF model was able to clearly distinguish between the differential and CA interactions, though it proved less effective in distinguishing between differential and CP interactions; see Figure 4b. The MS-only model can more proficiently resolve differential and CA interactions than the embeddings-only model, as shown in Figures 4c-d. This can be attributed to the fact that MINT [23] was trained on interactions present in STRING, where according to Figure 4a, it cannot properly separate differential and CA interactions. The weighted RF, MS-only, and embeddings-only models all proved less effective at distinguishing between differential and CP interactions; but, for all three models, their differential predictions were significantly greater than the CP predictions (Mann-Whitney U-test *p-value*<0.05). The outputs of two sequence-based PPI predictor models were also analyzed; see Figures 4e and 4f [17–18]. It is evident that PLM-interact cannot differentiate between the different interaction types; however, TUnA correctly ranks the CA interactions lowest but is unable to distinguish between differential and CP interactions.

## Discussion

Elucidating differential protein-protein interactions is instrumental for understanding condition-specific mechanisms of many cellular processes. As cancer impacts cell homeostasis, a deeper understanding of PPI rewiring events can be helpful in clinical contexts to identify novel putative targets. To this aim, our new machine learning model was developed to predict differential interactions between two experimental conditions, based on protein sequence and MS abundance data. This model was trained on eight distinct cell line-specific experimentally validated PPIs [4–6] and demonstrated proficiency in predicting differential interactions for both unseen proteins and conditions.

A weighted RF was used as the final model, as the data exhibited both label and cell line pair imbalances; see Table 2. The robustness of this weighted approach was revealed on the 14 external cell line pair test sets, as shown in Figure 3. There, in almost all cases, the weighted RF model outperformed its regular counterpart. This demonstrates that the weighting scheme enforced the model to learn a more generalizable decision boundary across cell line pairs when compared to the regular model, which was mostly influenced by the prevalent cell line pairs. Predicting differential interactions under unseen conditions proved a more challenging task, as compared to the sequence similarity-based split. This could be observed by the slightly degraded performance of our model in this task, relative to its performance on the sequence similarity-based split.

One limitation of our model is that though it can predict differential interactions between two conditions, it cannot specify in which condition the interaction is present. Also, though the model can specify if an interaction is conserved between conditions, it is not able to specify if the interaction is either present or absent in both conditions. Hence, the model predictions were further analyzed with respect to different conserved interaction types; see Figure 4. Evidently, the model can more easily distinguish between differential and CA interactions, as compared to distinguishing between differential and CP interactions. This can be attributed to the fact that most CP interactions are high confidence interactions based on STRING data, which may suggest that they are less likely to lose an interaction between two conditions; see Figure 4a. Another potential contributing factor is the training set for the sequence similarity-based split was composed of a disproportionate 20,938 CA and only 6,403 CP interactions. Unlike PLM-interact, TUnA ranked CA interactions lowest, which follows the pattern observed in the STRING combined scores; see Figures 4a, 4e and 4f. This suggests that sequence-wise CA PPIs are generally less likely to interact, relative to the other interaction types.

Additionally, characteristics of the different types of protein interactions were further analyzed. First, unique proteins that belong exclusively to a single category were considered, followed by enrichment analysis using enrichr [29] to detect significantly enriched gene ontology biological process (BP) terms [30]. Interestingly, CA interactions were mostly related to protein translation and ATP synthesis, as shown in Figure S1a. Conversely, differential interactions were enriched for multiple DNA processes and regulation of transcription; see Figure S1b. For CP interactions, no significant enrichment of BPs were found. Moreover, within protein complexes represented in the protein data bank, buried surface area and hydrophobicity of the protein-protein interfaces were investigated; see Figure S2 [31]. CA interactions were found to have the smallest buried surface area, suggesting that they generally do not form stable interactions. CP and differential interactions were determined to have similar surface areas. All three interactions exhibited interfaces with similar hydrophobicity levels; see Figure S3.

To our knowledge, no other current computational model that predicts differential PPIs between two experimental conditions exists. Our resulting final model is provided as an ensemble of four RFs that were trained on all available interactions, using different seeds. Predictions can be obtained using the scripts available on https://github.com/AramPapaz/Predicting-Differential-Protein-Protein-Interactions and represent the average predictions across the ensemble. We recommend focusing on the ranking of a certain prediction, rather than its raw value, as the RF models are uncalibrated. We believe that our model will prove advantageous in identifying dysregulated PPIs implicated in disease, such as cancer [32–33]. Ultimately, PPIs are involved at various stages of oncogenesis, including cell proliferation, survival, inflammation, invasion, and metastasis [34]. In summary, we intend that our model could serve as a selection criterion for experimental follow-up studies investigating deeper biochemical and molecular mechanisms attributing to rewired protein interactions.

## Data Availability

The ensemble of weighted RFs model can be downloaded from Zenodo (DOI: 10.5281/zenodo.23757927). The protein interactions and MS protein abundances can be retrieved from their respective studies [4–6, 19].

## Author Contributions

Aram Papazian: conceptualization, data analysis, analysis, model creation, and writing the draft for the manuscript. Volkhard Helms: conceptualization, discussion of results, edited the manuscript, and acquired funding.

## ACKNOWLEDGMENT

We would like to thank Dr. Michael Christopher Hutter and Anna Elizabeth Schmitz for editing the text and Nicholas Long Pham for helping to obtain MINT embeddings.

