## Supplementary for "Predicting Differential Protein-Protein Interactions"

### MS-Based Features

For the features having _**Zscore** as the suffix, the protein abundances were scaled using the modified z-score within each cell-line. The formula of the modified z-score is

$$z_{p,j}^{l} = \frac{0.6745 \times (x_{p,j}^{l} - median_k(x_{p,k}^{l}))}{MAD_p^l}$$

Where $x_{p,j}^{l}$ is the protein abundance of protein *p* in replicate *j* under condition *l*. $median_k(x_{p,k}^{j})$ is the median protein abundance of protein *p* across replicates *k* in condition *l*. $MAD_p^l$ is the median absolute deviation of protein *p*'s abundances from their median taken over the same replicates *k* in condition *l*.

For the JaccardNeighbors and ReciprocalTopK features, the Spearman correlation of each pair of proteins is computed based on their abundance values. This is calculated for all proteins measured in the proteome. Then, for each protein, only the top 25% most correlated partners are kept. To compute these features between two proteins, the following formulas were used:

$$JaccardNeighbors(p_i, p_j) = \begin{cases} \frac{|N_i \cap N_j|}{|N_i \cup N_j|} & if\ |N_i \cup N_j| > 0 \\ 0 & otherwise \end{cases}$$

$$ReciprocalTopK(p_i, p_j) = \begin{cases} 1 & if\ p_i \in N_j, p_j \in N_i \\ 0 & otherwise \end{cases}$$

Where $p_i$and $p_j$ represent proteins *i* and *j,* respectively. $N_i$ and $N_j$ represent the set of top 25% most correlated partners of proteins *i* and *j,* respectively.

### PPI Properties

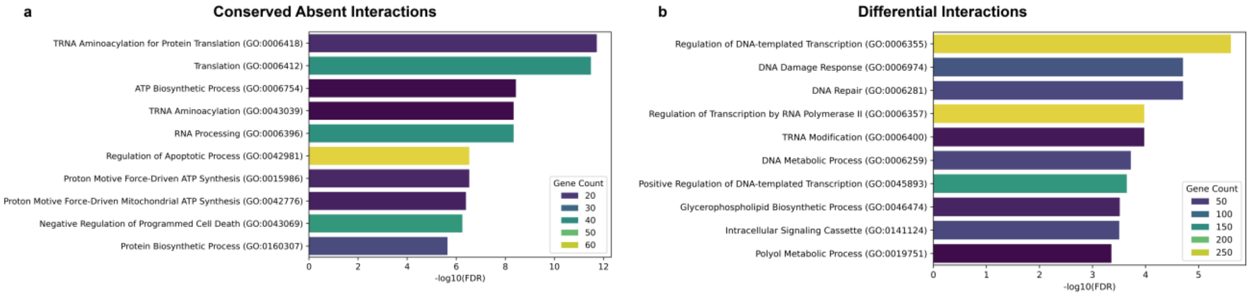


Figure S1. Significantly enriched gene ontology biological processes for Conserved_Absent (**a**) and differential (**b**) interactions

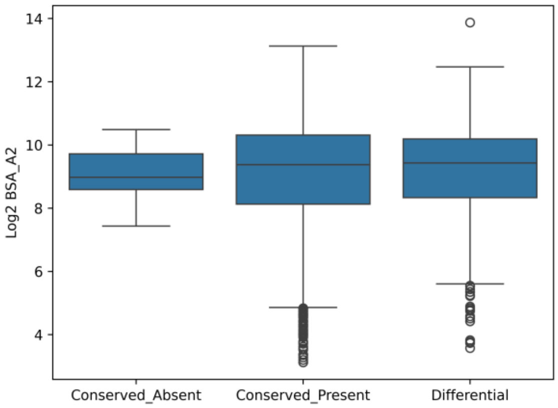


Figure S2. Buried surface areas of protein-protein interfaces for the different types of interactions. These only include the 2,077 PPIs that had protein databank entries.

Concerning the buried surface area between two interacting proteins, we first searched for protein data bank (PDB) entries that include both proteins. Then, only the ones where the closest pair of atoms belong to protein 1 and 2, respectively, are at most 5 Ångstroms away from each other were kept. This threshold assumes they are close enough to interact. If two proteins were part of many PDB entries that met the distance criteria, the entry that had the best resolution was kept. Then the buried surface area was computed using:

$$\frac{SASA_1 + SASA_2 - SASA_{1,2}}{2}$$

Where *SASA* represents the solvent-accessible surface area for a certain protein.

To compute hydrophobicity, all residues in the PDB entry that lost at least 1 Ångstrom of exposed area upon binding were defined as interface residues. Each interface residue was assigned its Kyte-Doolittle hydropathy score and the final hydrophobicity level was calculated by taking the average of the interface residue scores.

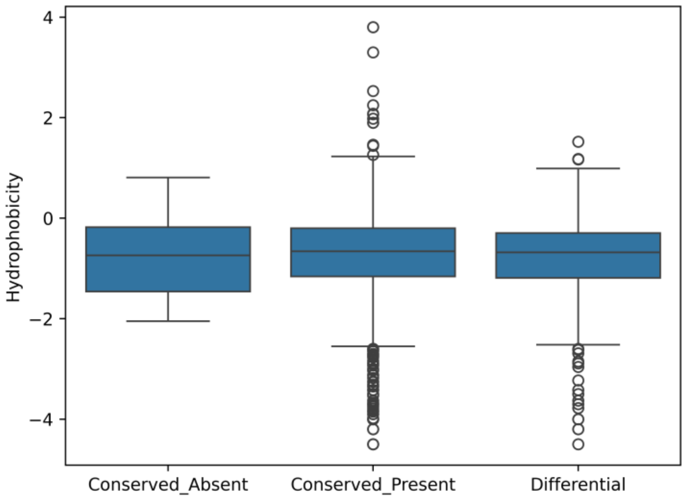

Figure S3. Hydrophobicity levels of protein-protein interfaces for the different types of interactions. These only include the 2,077 PPIs that had protein databank entries.
